# The generation of persister cells is regulated at the initiation of translation by (p)ppGpp

**DOI:** 10.1101/2020.09.17.300954

**Authors:** Roberto C. Molina-Quiroz, Andrew Camilli

**Author notes:** Address correspondence to Roberto Molina-Quiroz.

## Abstract

Bacterial persistence is a non-heritable phenotypic trait characterized by a dormant state that leads to tolerance to different antibiotics. Several mechanisms contributing to persister cells generation have been identified. Among these, is the signaling molecule (p)ppGpp, but knowledge of how this molecule regulates persister generation is incomplete. Here, we show an increase of the persister fraction of uropathogenic *Escherichia coli* (UPEC) that correlates with the time of protein synthesis inhibition and a decrease in the availability of antibiotic target. Specifically, the arrest of translation initiation induces bacterial survival to ampicillin and ciprofloxacin in a (p)ppGpp-dependent manner. These findings support a global mechanism of persister cell generation and establish a regulatory role of the (p)ppGpp molecule in this phenomenon.

**Importance:** The study of persister cell formation is relevant because this bacterial subpopulation is involved in the emergence of antibiotic resistance and the generation of chronic infections. A role of the (p)ppGpp molecule in the generation of the persister fraction has been described, but the identification of the regulatory mechanism mediated by this alarmone during protein translation and its contribution to persistence has not been described to date. In this work, we show that (p)ppGpp regulates the generation of persister cells at the initiation of the protein synthesis process in UPEC. Our results also suggest that a (p)ppGpp-dependent regulation of translation, might be a global mechanism for the generation of the persister fraction.

## Introduction

Bacterial persistence is a transitory, non-genetically heritable dormant state that leads to tolerance to normally lethal concentrations of different antibiotics in several bacterial species. After antibiotic challenge, this fraction can resume growth when stress is relieved. Persister cells play roles in virulence, relapsing infections, and facilitate evolution of antibiotic resistance (1). Different mechanisms and signaling molecules involved in persister cell generation have been described (1). Among these are inhibition of translation (2), cyclic AMP (cAMP) (3), and the (p)ppGpp alarmone molecule (4, 5). The regulatory effects of (p)ppGpp on bacterial growth, transcription and DNA replication have been widely studied (6). However, a deeper understanding of (p)ppGpp regulation of translation and its impact on persistence is lacking. Here we show that the availability of antibiotic-target molecule is critical for the generation of persister cells, and that this phenomenon is tightly regulated at the level of initiation of protein synthesis in a (p)ppGpp-dependent fashion.

## Results and Discussion

A role for (p)ppGpp in persistence has been widely proposed, yet a deeper understanding of (p)ppGpp regulation of translation and its impact on persistence is still lacking (4, 7, 8). Aiming to explore the role of (p)ppGpp in persistence of UPEC, we pre-exposed exponentially growing cells for 30 min to DL-Serine hydroxamate (SHX), an inhibitior of Seryl-tRNA synthetase (SerS). SHX-treated and control cultures where then exposed to high concentrations of ampicillin or ciprofloxacin for 4 hours to select the persister fraction. SHX has been widely used to induce the persister state through the arrest of protein synthesis generated by the increase in the (p)ppGpp level, the so called stringent response (5, 9). As expected, an increased survival to both ampicillin and ciprofloxacin was observed in cultures pre-exposed to SHX (Figure 1A). The role of (p)ppGpp in this phenomenon was assessed using a Δ*relA* Δ*spoT* double mutant, which is unable to synthetize (p)ppGpp, *i.e*., is (p)ppGpp^0^. The toxic effect generated by SHX in cultures of the (p)ppGpp^0^ (compared to wt), might be explained by a global change in the gene expression profile mediated by SHX. A decreased ability to either upregulate and/or downregulate genes related to metabolism, translation, nucleic acid metabolism, transport and others has been shown when a (p)ppGpp^0^ strain is challenged with SHX (10).

**Figure 1.**
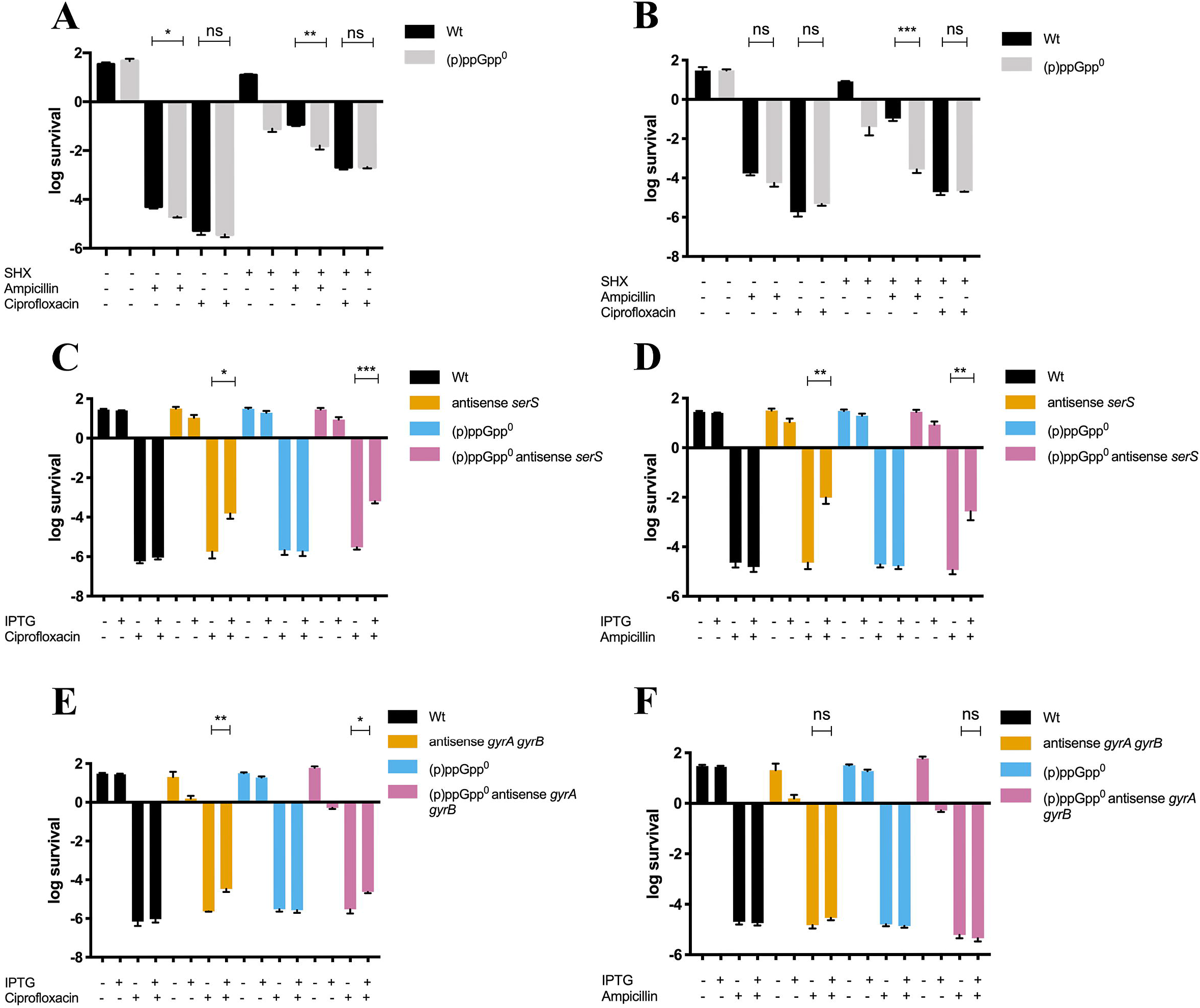
Availability of the antibiotic target is required for the generation of persister cells. Bacterial survival to ampicillin or ciprofloxacin was addressed by CFU counting after induction of stringent response by incubation of cultures in the presence of SHX (A). Persister cells generation was assessed in cultures exposed to ampicillin (B) or ciprofloxacin (C) in cultures that had been previously grown in the presence or absence of IPTG to decrease the expression of *serS* by anti-sense RNA induction. The ability to generate persister cells was addressed by CFU counts in cultures pre-exposed 3 min to SHX (D). The expression of *gyrA* and *gyrB* was downregulated by generation of an IPTG-dependent antisense RNA and the cultures were subsequently challenged with ciprofloxacin (E) or ampicillin (F) to evaluate the generation of persister cells by CFU counting. Graphs represent the average of at least three independent assays. Statistical significance was calculated using Student’s two-tailed t-test (*p<0.05, **p<0.01, ***p<0.001, ****p<0.0001).

When cultures were pre-exposed to SHX and then challenged with ampicillin, a 10-fold decrease in bacterial survival was observed in the (p)ppGpp^0^ strain compared to the wt, suggesting a role of (p)ppGpp in the generation of persister cells (Figure 1A). However, this phenomenon did not occur in cultures treated with ciprofloxacin (Figure 1A), suggesting a (p)ppGpp-independent mechanism of persistence to ciprofloxacin. SHX induces the stringent response by inhibition of Seryl-tRNA synthetase (9), but it is possible that a 30 min pre-exposure may have other unknown effects on cell physiology that are not necessarily related to the stringent response (11). We did two experiments to examine this. First, we tested if a shorter pre-exposure to SHX, which should still induce the stringent response and arrest translation, impacts persister cell generation. Indeed, just 3 min of SHX pre-exposure resulted in increased survival to both antibiotics, though not to the extent observed after 30 min of pre-exposure (Figure 1B). Again, a (p)ppGpp-dependent effect on survival was observed for ampicillin but not ciprofloxacin. Second, we tested the role of (p)ppGpp in persistence by an independent method, which was to genetically induce the repression of *serS* via an antisense RNA. To accomplish this, we inserted the IPTG-inducible Ptac promoter downstream of *serS* in the antisense orientation. This genetic tool has been used and validated previously to downregulate the expression of genes related to persistence (12). Also, to evaluate that a decreased production of Seryl-tRNA synthetase generated as a consequence of the downregulation of *serS*, induces arrest of growth in both genetic backgrounds (wt and (p)ppGpp^0^), we plated the strains on LB and LB supplemented with IPTG agar plates. After overnight growth at 37°C no growth was observed for strains containing the Ptac promoter LB/IPTG plates (not shown). These results show that induction of antisense RNA inhibits the expression of *serS* leading to arrest of bacterial growth. Downregulation of *serS* increased bacterial survival to ampicillin and ciprofloxacin in both wt and (p)ppGpp^0^ backgrounds (Figure 1B and C). These results suggest that global inhibition of protein synthesis, and likely the inability to synthetize antibiotic-target proteins, might lead to persistence. There was a difference in the role of (p)ppGpp in survival to ciprofloxacin depending on whether SHX or antisense repression of *serS* was used to induce the stringent response. This could be explained by possible unknown effects of SHX on DNA replication or other processes contributing to persistence. This difference notwithstanding, our results support the existence of (p)ppGpp-dependent and (p)ppGpp-independent mechanisms for regulation of persistence when either the activity of SerS or its expression are inhibited. A 4-fold difference in survival was observed between wt and (p)ppGpp^0^ cultures exposed to ampicillin (Fig 1A). However, non-significant differences were determined later for the same experimental condition (Fig. 1B and D respectively). This might be explained by the high variability of the persister assay itself.

We hypothesized that the formation of persister cells observed upon inhibition of SerS activity or expression may be due in part to the arrest of protein synthesis resulting in reduced availability of the antibiotic targets themselves. To explore this, we constructed a strain with P_tac_-mediated antisense repression of *gyrA* and *gyrB*, which encode the DNA gyrase target of ciprofloxacin (13). Similarly as described above for the antisense *serS* strain, no colonies were observed after overnight growth of the antisense *gyrA/gyrB* strains in both wt and (p)ppGpp^0^ backgrounds on LB plus IPTG agar plates (not shown). This suggests that a decreased amount of DNA gyrase generated as a consequence of the downregulation of its cognate genes, leads to arrest of bacterial growth. IPTG-induced downregulation of *gyrA* and *gyrB* prior to the antibiotic challenge resulted in an increased survival to ciprofloxacin in the wt and (p)ppGpp^0^ backgrounds (Figure 1E), but not to ampicillin (Figure 1F). These results suggest that a decreased availability in the antibiotic target molecule leads to the generation of persister cells.

Our results suggest that limitation of target protein amounts can reproduce the phenotype of persistence that is triggered by antisense repression of *serS*, and that this phenomenon is not dependent on (p)ppGpp. Our results are further supported by recent findings showing that (p)ppGpp modulates DNA replication by regulating the expression of gyrase (14). We chose not to pursue a similar strategy for ampicillin because *E. coli* has eight Penicillin Binding Protein (PBP) targets (15). Nevertheless, previous findings show that (p)ppGpp regulates ampicillin persistence via a shutoff of peptidoglycan biosynthesis during diauxic shifts (16). These results suggest that persistence to cell wall acting agents might be regulated by (p)ppGpp by decreasing the levels of antibiotic-target molecules.

Based on our results and those of other groups mentioned above, we reasoned that to survive to antibiotic challenge through persister cell formation, bacteria must regulate the level of antibiotic target proteins by fine tuning transcription and/or translation, and (p)ppGpp might play a role. We tested this idea using different inhibitors that specifically affect either transcription or different stages of protein synthesis. Cultures of wt and (p)ppGpp^0^ strains were pre-exposed for 30 min to rifampicin to inhibit transcription, and subsequently challenged with ampicillin or ciprofloxacin. Both strains with halted transcription exhibited increased survival to ampicillin and ciprofloxacin when compared to the untreated controls, as previously reported (2). However, no contribution of (p)ppGpp on persistence was observed (Figure 2A). Similarly, when cultures were pre-exposed for 30 min to puromycin, chloramphenicol, or tetracycline to target translation at different stages (17), an increased survival for ampicillin and ciprofloxacin was observed in both genetic backgrounds (Figure 2B-D). These results indicate that (p)ppGpp has no effect on the generation of persister cells when protein synthesis is arrested by premature release of the nascent polypeptide chain (puromycin) (17), prevention of the peptide bond formation (chloramphenicol) (17), or prevention of the stable binding of the EF-Tu-tRNA-GTP complex to the ribosome (tetracycline) (17). In contrast, when translation was arrested by blocking the release of EF-Tu from the ribosome with kirromycin (17), a 10-fold decrease in the persister fraction in the (p)ppGpp^0^ background was observed in cultures exposed to ciprofloxacin but not to ampicillin. These results suggest that by affecting the EF-Tu-ribosome interaction (p)ppGpp generates an antibiotic-specific response leading to the generation of persister cells (Figure 2E). In addition, a ~100 and 10-fold decrease in the survival to ampicillin and ciprofloxacin, respectively, of the (p)ppGpp^0^ strain compared to wt was observed in cultures that were pre-treated with kasugamycin (Figure 2F). This antibiotic inhibits the translation initiation of canonical (but not leaderless) mRNAs by preventing the stable interaction of the initiator tRNA with the start codon (17, 18). Altogether, these results demonstrate that the availability of antibiotic target molecules is required for the generation of the persister fraction, and that this phenomenon is finely regulated at the initiation of translation level by (p)ppGpp. Regarding the (p)ppGpp-independent mechanism, a decrease in ATP levels, proton gradient regulation and other processes have been implied in bacterial persistence (1).

**Figure 2.**
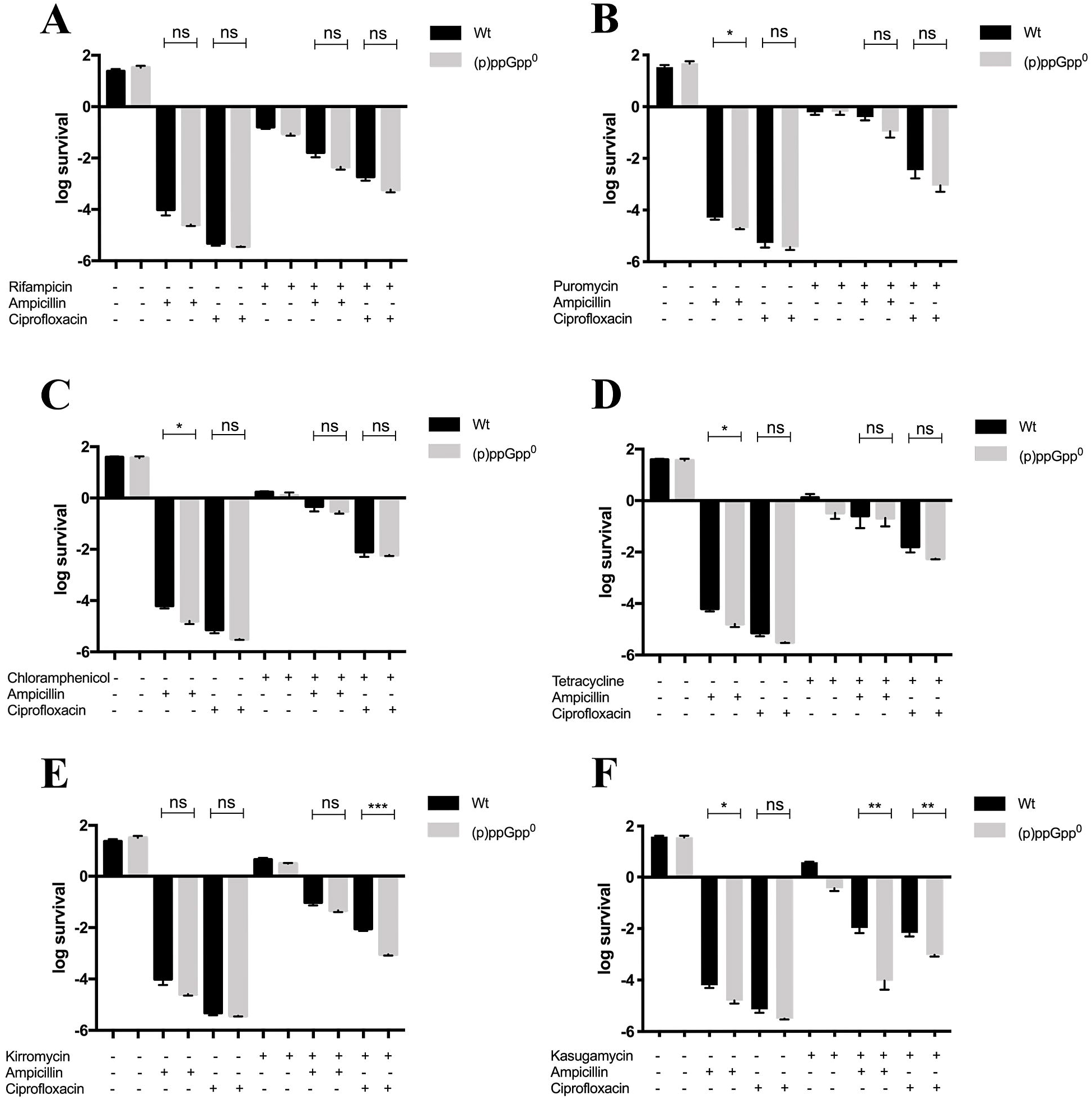
The initiation of protein synthesis is a global mechanism of persister cell generation that is regulated by (p)ppGpp. The generation of persister cells was addressed by CFU counting in cultures pre-treated with different inhibitors of transcription or translation, and subsequently challenged with ampicillin or ciprofloxacin. Transcription was inhibited by rifampicin (A), whereas different stages of translation were inhibited by puromycin (B), chloramphenicol (C), tetracycline (D), kirromycin (E) or kasugamycin (F). Graphs represent the average of at least three independent assays. Statistical significance was calculated using Student’s two-tailed t-test (*p<0.05, **p<0.01, ***p<0.001, ****p<0.0001).

Our findings are supported by a recent report showing that (p)ppGpp inhibits the activation of IF2 leading to the attenuation of initiation of translation during the switch from active growth to quiescence in the Gram-positive bacterium *Bacillus subtilis* (19). Our work in a Gram-negative (UPEC) suggests for the first time that the regulation of the initiation of protein synthesis mediated by (p)ppGpp might be a common evolutionary trait in bacteria, leading to the survival to harsh conditions such as antibiotic and nutritional stress.

## Materials and Methods

### Bacterial strains and growth conditions

Cultures of UPEC CFT073 were grown with shaking at 37°C in LB Miller (hereafter LB). LB was supplemented with 0.13 μg/ml ciprofloxacin, 1250 μg/ml ampicillin, 500 μg/ml SHX, 6.3 μg/ml tetracycline, 6 μg/ml chloramphenicol, puromycin 100 μg/ml, rifampicin 20 μg/ml, 100 μg/ml kirromycin, or 750 μg/ml kasugamycin when required. Induction of antisense RNA was conducted by addition of 100 μM IPTG and incubation at 37°C for 90 min until cultures reached OD_600_=0.2.

(p)ppGpp^0^ (Δ*rel4*::FRT/Δ*spoT*::FRT) and antisense strains were constructed by recombination of PCR products and backcrossed to the wt genetic background as described (3). All mutants used in this work were subjected to whole genome sequencing and variant analyses to check for undesired off target or spontaneous mutations.

### Persister assays

Overnight cultures were diluted 1000-fold in fresh LB and incubated at 37°C with aeration until OD_600_=0.1. Then, transcription or translation inhibitory drugs were added and incubated at 37°C with aeration for 30 min (typically reaching ~2 x 10^8^ CFU/ml). Cultures were subsequently challenged independently for 4 h with ampicillin or ciprofloxacin at 37°C with aeration. Aliquots were washed and plated on LB agar plates for CFU counting after overnight incubation at 37°C. Survival was determined by dividing the number of CFU/ml in the culture after 4 h of exposure to antibiotics by the number of CFU/ml before adding the antibiotic.

## Acknowledgments

We thank David Lazinski for scientific discussions and Cecilia Silva-Valenzuela for critical reading of this manuscript and scientific feedback. This work was supported by Fondecyt Iniciación en Investigación 11190158 (RCM-Q). Centro de Estudios Científicos (CECs) is funded by the Centers of Excellence Basal Financing Program of CONICYT PB-01.

## Author Contributions

RCM-Q performed experiments. RCM-Q, and A.C. designed experiments, wrote the manuscript, provided materials and strains. Authors discussed the results and commented on the manuscript.

